# Synaptic Toxicity of OGA Inhibitors and the Failure of Ceperognastat

**DOI:** 10.1101/2025.05.09.648606

**Authors:** Jonathan Meade, Haylee Mesa, Lei Liu, Qi Zhang

## Abstract

O-GlcNAcase inhibitors (OGAi) have been proposed as therapeutics for Alzheimer’s disease due to their ability to increase O-GlcNAcylation of tau and reduce its aggregation. However, the recent failure of ceperognastat in a Phase II trial—marked by accelerated cognitive decline in the treatment arm—has raised concerns about the safety of this therapeutic class. Here, we evaluated the acute synaptic effects of three structurally distinct OGAi (ceperognastat, ASN90, and MK8719) in mouse hippocampal slices. Electrophysiological recordings revealed that all three compounds impaired both short- and long-term synaptic plasticity, as evidenced by reduced paired-pulse facilitation/depression and suppressed long-term potentiation.

Immunohistochemistry showed altered synaptic protein levels, with increased PSD-95 and reduced Synaptophysin 1 in neurons, alongside a biphasic shift in Tau phosphorylation. These findings indicate that OGAi produce rapid and convergent synaptotoxic effects across pre- and postsynaptic compartments, likely reflecting a class-wide mechanism. We argue that electrophysiological screening should be standard in CNS drug development and caution against targeting essential synaptic processes in chronic neurodegenerative conditions.

## Introduction

Adding O-linked β-N-acetylglucosamine (O-GlcNAcylation) to serine or threonine residues is a reversible post-translational modification in many proteins^1^. This process is regulated by the reciprocal actions of O-GlcNAc transferase (OGT, adding O-GlcNAc) and O-GlcNAcase (OGA, removing O-GlcNAc). Neuronal proteins essential for synaptic transmission and plasticity— including synapsin and AMPA receptors—undergo O-GlcNAcylation. Genetic ablation of OGA causes perinatal lethality, and heterozygous OGA knockout mice exhibit deficits in both long-term potentiation (LTP) and long-term depression (LTD), underscoring the critical role of dynamic O-GlcNAc cycling in neural function^2^. Despite this, OGA became a therapeutic target based on its ability to modulate tau pathology in animal models: O-GlcNAcylation competes with tau phosphorylation, and preclinical studies reported that OGA inhibition reduces tau aggregation^3^. These findings led to the development of OGA inhibitors (OGAi) for Alzheimer’s disease and other tauopathies. However, in the Phase II PROSPECT-ALZ trial of the OGAi ceperognastat□, participants receiving the 3 mg dose exhibited significantly faster cognitive decline than placebo across multiple cognitive measures, raising serious concerns about drug-induced neurotoxicity□. This study evaluates whether preclinical synaptic deficits induced by OGAi could have anticipated these adverse outcomes, and whether such effects reflect a class-wide liability for OGA-targeting therapeutics.

## Methods

Ceperognastat (LY3372689), Egalognastat (ASN90), and MK8719, were purchased from MedCehm Express. 300-μm mouse hippocampal slices were treated with DMSO or OGAi for ∼ 4 hours for whole-cell patch-clamp recording of CA1 neurons. Slices were subsequently used for immunohistochemistry. Methods are detailed in supplementary information.

## Results

We assessed short-term synaptic plasticity by measuring paired-pulse facilitation (PPF) and depression (PPD) in mouse hippocampal slices treated with 10□μM OGAi or DMSO for 4 hours. PPF was defined as the response increase across five pulses in the first stimulation episode (Figure 1A1), and PPD as the reduction in the first response across episodes (Figure 1A2). All three OGAi reduced both PPF and PPD compared to control (Figure 1B), indicating impaired presynaptic plasticity. Next, we evaluated long-term potentiation (LTP) and found that all OGAi significantly suppressed LTP induction (Figure 1C). These results demonstrate that OGAi from three distinct pipelines disrupt both short- and long-term synaptic plasticity, implying pre- and postsynaptic alternation.

**Figure 1.**
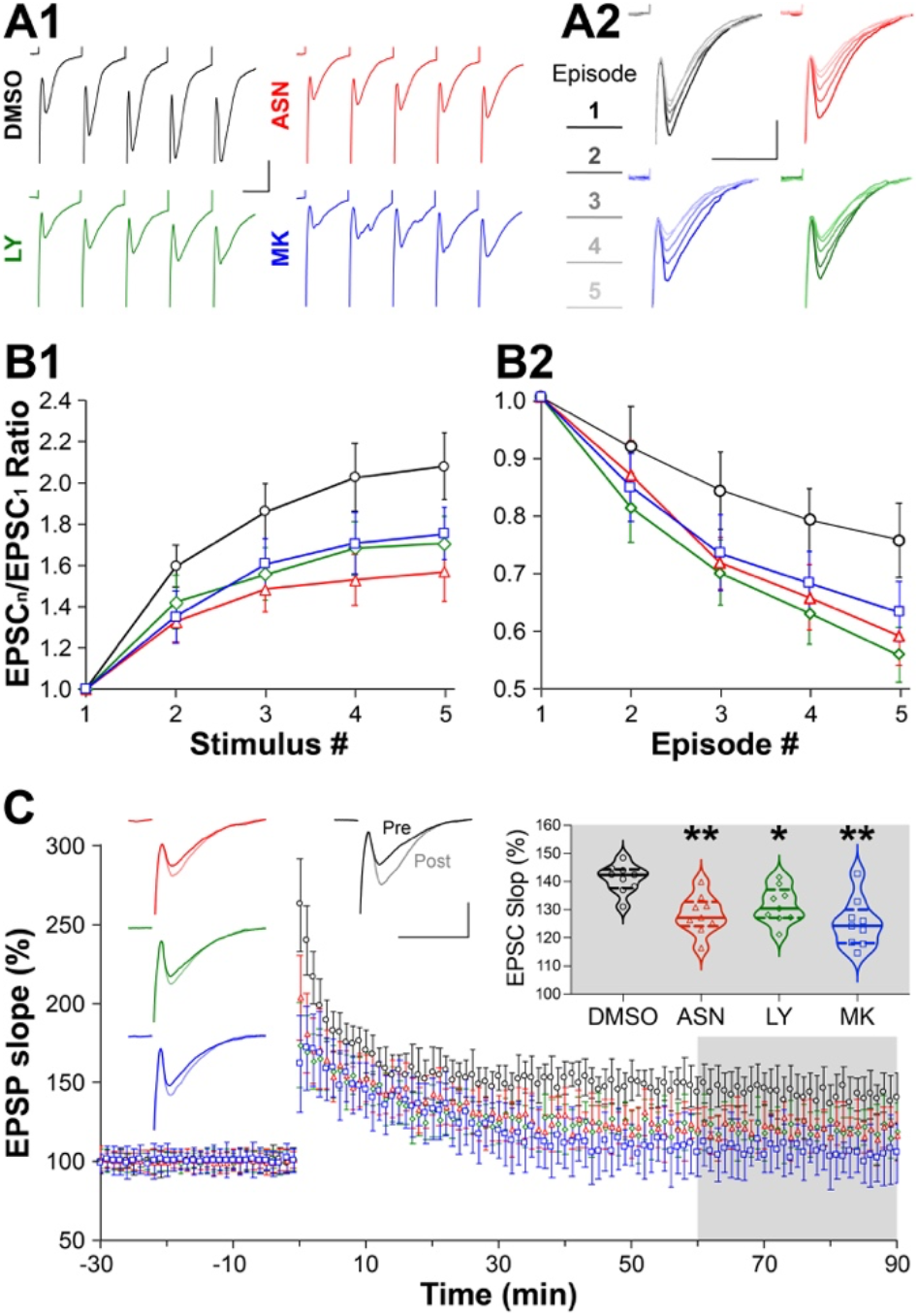
OGA inhibitors alter synaptic plasticity. **A**, sample traces of whole-cell patch clamp recording of hippocampal CA1 pyramidal neurons during the first episode of paired-pulse stimulation (**A1**) and aligned first response from every episode (**A2**). Scale bar, 20ms (horizontal) and 0.5pA (vertical). The color coding for DMSO (black), ASN90 (i.e., ASN, red), LY3372689 (i.e., LY, green), and MK8719 (i.e., MK, blue) is used in all plots. **B**, plots of EPSC ratio for the responses during the first episode (**B1**) and the first responses of five episodes (**B2**). **C**, LTP measured as relative EPSP slopes after the treatments of OGAi or DMSO. Left insets are sample traces pre- and post 100-Hz 1-s tetanus stimulation. Right inset is a violin plot of the average values during the last 30-s recording (i.e., gray box in the plot). Welch’s ANOVA test, W (DFn, DFd) = 10.61 (3.000, 17.50), *p* = 0.0003; Kolmogorov-Smimov test, *p* = 0.0063 (ASN *vs*. DMSO), 0.0336 (LY *vs*. DMSO), and 0.0063 (MK *vs*. DMSO). N = 3 mice and n = 3 slices (i.e., 9 recordings in total for every treatment).

To further assess synaptic changes following acute OGAi treatment, we performed immunohistochemistry using well-validated antibodies (Tables S1 and S2). We observed a significant increase in PSD-95 immunolabeling within Tuj1-positive/GFAP-negative regions (Figure 2A & C). Additionally, OGAi reduced Synaptophysin 1 labeling in Tau-positive neurites (Figure 2B & D). Together, these findings demonstrate that OGAi induces morphological changes in both dendritic and synaptic compartments. These structural alterations were accompanied by a biphasic shift in pTau/Tau ratios (Figure 2E), confirming on-target pharmacodynamic engagement.

**Figure 2.**
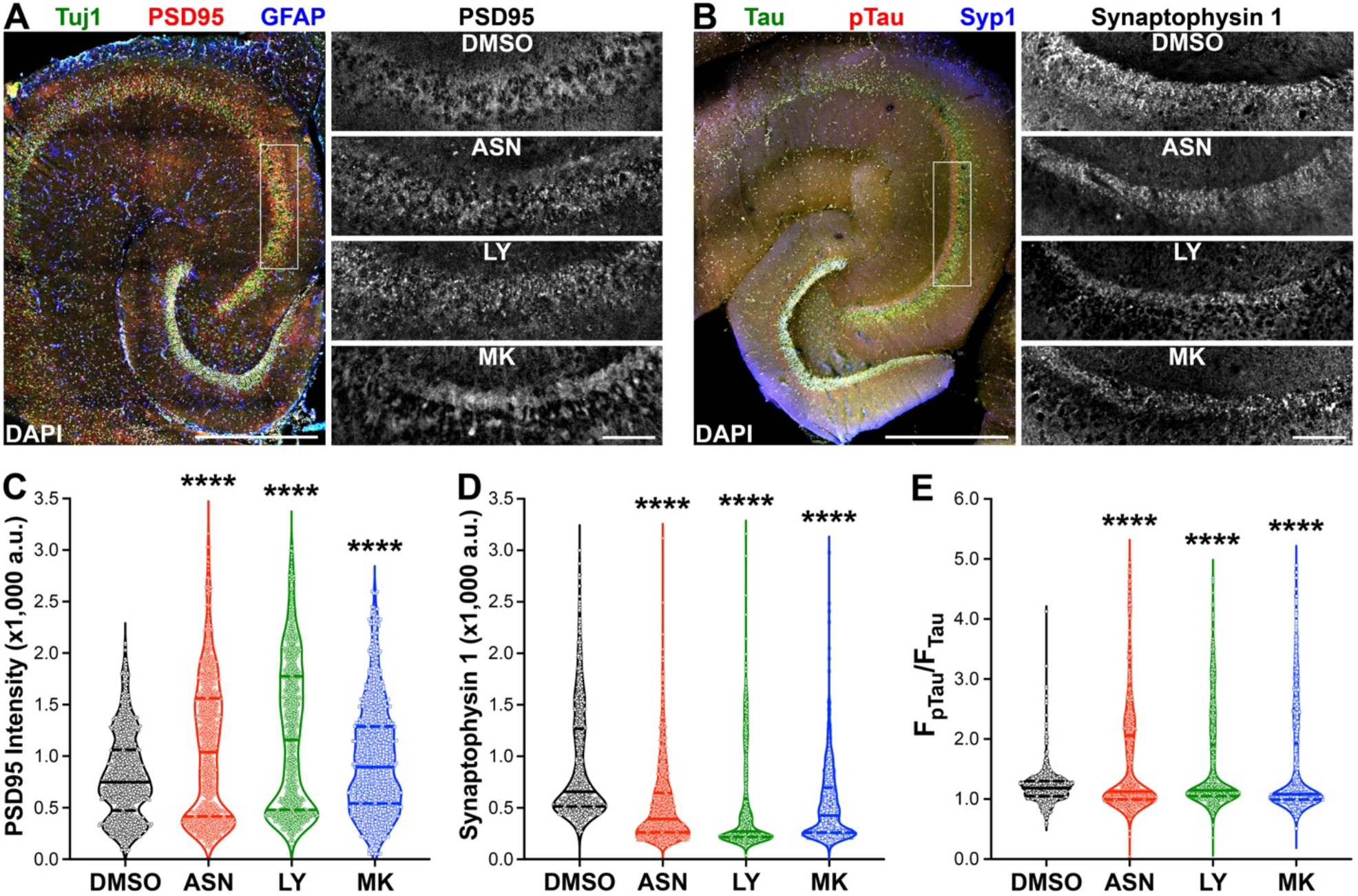
OGA inhibitors alter synaptic protein levels and Tau phosphorylation. **A**, sample images of fluorescence immunocytochemistry for PSD95, Tuj1, GFAP, and DAPI. Scale bar, 0.5 mm. Images on the left are zoom-in images of PSD95 immunofluorescence in the cell-body layer (indicated by the white box) for all four treatments. Scale bar, 50 μm **B**, sample images of fluorescence immunocytochemistry for Synaptophysin 1, Tau, pTau, and DAPI. Images on the left are zoom-in images of Synaptophysin 1 immunofluorescence in the cell-body layer (indicated by the white box) for all four treatments. **C&D**, violin plots of PSD95 (**C**) and Synaptophysin 1 (**D**) immunofluorescence intensities. **E**, violin plots of pTau *vs*. Tau ratios from the same ROIs of Synaptophysin 1. Welch’s ANOVA test for PSD95, W (DFn, DFd) = 90.96 (3.000, 2168), *p* < 0.0001; for Synaptophysin 1, W (DFn, DFd) = 153.30 (3.000, 2187), *p* < 0.0001; for pTau/Tau, W (DFn, DFd) = 128.40 (3.000, 1918), *p* < 0.0001. Kolmogorov-Smimov test for all three, *p* < 0.0001 (ASN *vs*. DMSO, LY *vs*. DMSO, and MK *vs*. DMSO). N = 9 slices from 3 mice (3 ea), n = 999 ROIs (111 randomly selected from every slice).

## Discussion

Ceperognastat’s failure echoes the repeated clinical disappointments of BACE inhibitors (BACEi), shown to impair cognitive function through acute disruption of neurotransmission^6^. Although mechanistically distinct, both drug classes share a common outcome: BACEi impair synaptic function by prolonging the half-life of cleavable synaptic substrates, while OGAi disrupt synaptic integrity by stagnating synaptic protein O-GlcNAcylation. Our findings serve as a cautionary tale for ongoing OGAi programs and future drug development efforts targeting chronic cognitive or neurological disorders. We strongly advocate for the routine incorporation of electrophysiological assessments into toxicology panels to identify synaptic liabilities before advancing compounds into clinical trials. In this study, we investigated the synaptic consequences of OGAi administration, reportedly achieving > 95% OGA enzyme occupancy^4^— levels that in hindsight reflect a fundamentally flawed therapeutic strategy for a treatment of chronic neurological cognitive diseases. Further mechanistic study is needed to understand the molecular underpinnings of this drug-induced cognitive decline and to prevent similar adverse outcomes in future clinical programs.

## Acknowledgments

We thank Drs. Hideki Iwamoto for helping with adult brain slice preparation.

## Additional Information

All experimental procedures are described in detail in the Supplementary Information.

