## Supplemental Info for "Synaptic Toxicity of OGA Inhibitors and the Failure of Ceperognastat"

MGB research building, BLND452, 65 Landsdowne St, Cambridge, MA, 02139, USA

5353 Parkside Dr., Florida Atlantic University, Jupiter, FL 33458, USA.

**Table S1: Primary antibodies used for immunocytochemistry**

| **Combo** | **Antigen** | **Species** | **Type** | **Supplier** | **Cat. No.** | **Dilution** |
| --- | --- | --- | --- | --- | --- | --- |
| **1** | GFAP | guinea pig | polyclonal | SYSY | 173004 | 1:250 |
|  | Tuj1 | rabbit | polyclonal | Sigma | T3952 | 1:200 |
|  | PSD95 | mouse | monoclonal | Antibodies Inc | 75-028 | 1:500 |
| **2** | Synaptophysin1 | guinea pig | polyclonal | SYSY | 101004 | 1:500 |
|  | Tau | chicken | polyclonal | Aveslabs | AB_2313563 | 1:2000 |
|  | pTau | mouse | monoclonal | Thermo Scientific | AT8 | 1:400 |

**Table S2: Secondary antibodies used for immunocytochemistry**

| **Antigen** | **Species** | **Fluorophore** | **Supplier** | **Cat. No.** | **Dilution** |
| --- | --- | --- | --- | --- | --- |
| Chicken IgY (H+L) | goat | CF®488A | Biotium | 20020 | 1:1000 |
| Guinea pig IgG (H+L) | goat | CF®647 | Biotium | 20041 | 1:1000 |
| Mouse IgG (H+L) | goat | CF®568 | Biotium | 20101 | 1:1000 |
| Rabbit IgG (H+L) | goat | CF®488A | Biotium | 20019 | 1:1000 |

**Methods**

***Brain slice preparation***

All chemical and biological reagents, if not specified, were purchased from Sigma-Aldrich. All three OGA inhibitors (OGAi) were stored as 10 mM DMSO solution in -80°C freezer. Animal use was approved by FAU IACUC (A23-26). All 6-month-old C57B6/J male mice were purchased from the Jackson Laboratory and husbanded in FAU vivarium until use. 300-μm thick coronal slices containing hippocampal formations were prepared using Leica VT1000S and incubated in recovering artificial cerebrospinal fluid (aCSF, in mM: 92 NaCl, 2.5 KCl, 1.25 NaH_2_PO_4_, 30 NaHCO_3_, 20 HEPES, 25 glucose, 2 thiourea, 5 Na-ascorbate, 3 Na-pyruvate, 2 CaCl_2_·4H_2_O and 2 MgSO_4_·7H_2_O). During the recovery, DMSO (1:1000 dilution) or 10 μM OGAi (1:1000 dilution from the stock solutions) were applied to the recovery aCSF for ~4 hours.

***Electrophysiology***

All following tests and analyses were done blind. Whole-cell patch-clamp recording of CA1 neurons was performed in the recording aCSF (in mM: 119 NaCl, 2.5 KCl, 1.25 NaH_2_PO_4_, 24 NaHCO_3_, 12.5 glucose, 2 CaCl_2_·4H_2_O and 2 MgSO_4_·7H_2_O) with the same concentrations of DMSO or OGAi as those during recovery. Schaffer collateral stimulations for paired-pulse facilitation and depression (30Hz 0.15s, i.e., 5 pulses per episode and five episode with 10-second interval in between) and long-term potentiation (100-Hz 1-second tetanus stimulation) were delivered via a micro-electrode. For EPSC, membrane potentials were held between -70 and -40 mV). All experiments were performed using ROE-200 dual-manipulator and MPC-200 controller (Sutter Instruments), MultiClamp 700B and DigiData 1440A controlled by pCLAMP 10 (Molecular Devices). Clampfit 10.6 was used for data analysis and extraction.

***Immunocytochemistry***

After recording, slices were fixed using 4% paraformaldehyde in phosphate-buffered saline (PBS, in g for 1L, 8 NaCl, 0.2 KCl, 1.15 Na_2_HPO_4_•7H_2_O, 0.2 KH_2_PO_4_, pH = 7.35) for 1 hours at room temperature. The fixation and all subsequent steps were carried out in 12-well plates and on a plate shaker set at 60 rpm. After three times wash with PBS, slices were permeabilized with 3% Triton X-100 (in PBS) for 1 hour and blocked by 5% Goat serum (Abcam), 5% (W/V) protease-free BSA (W/V) and 0.25% Triton X-100 (in PBS) for 1 hours at room temperature. Primary antibody combos (Table S1) were prepared in fresh block solution and incubated with those slices at 4°C overnight. After the removal of primary antibodies, the slices were washed three times with PBS containing 5% Goat serum, 0.5% (W/V) protease-free BSA (W/V) and 0.25% Triton X-100. The secondary antibody combos (Table S2) were applied in the same solution for 4 hours at room temperature. After the removal of secondary antibody, slices were stained with Hoechst 33342 nucleic acid stain (Invitrogen) for 30 minutes at room temperature and washed with PBS and Milli-Q water twice for each. All slices were mounted in VECTASHIELD® antifade mounting media (Vector Laboratories).

***Image acquisition and analyses***

Multichannel confocal images of those brain slices were acquired using a Nikon A1R laser scanning confocal microscope equipped with a Nikon Plan Apo VC 20X N.A. 0.75 objective and controlled by NIS-Elements AR (Version 5.42.04). The acquisition settings are as follows: DAPI (i.e., Hoechst 33342), 405nm for excitation, 430-475nm for emission, 1.0 for laser power, 60 for gain, 17.88μm pinhole size; CF488A®, 488nm, 500-550nm, 0.3, 80, 26.82μm; CF568®, 560nm, 570-616nm, 3.0, 80, 26.82μm; CF647®, 641nm, 660-710nm, 5.0, 90, 26.82μm. With such settings, the pixel intensities corresponding to biological specimens never exceeded the dynamic range of 10-bit image (i.e., 0 – 4,095). Z-stack step size is 3 μm.

All images were imported into FIJI (ImageJ2) by Bio-Formats (plug-in). For every brain slice, three Z-consecutive sections in the middle of the image stack were Z-projected using their average pixel intensities. Hippocampal formations were selected from the project images. Four cell-free regions of interest (ROIs) were selected, and the mean as well as the standard deviation of their pixel intensities for every channel were calculated. ROI selection in individual channel was done by thresholding, which was set at 3 times of standard deviation above the mean background pixel intensities. For postsynaptic PSD95, Tuj1-positive, GFAP- and DAPI-negative ROIs were generated via Boolean operations of all three corresponding images. Similar operations were done to generate ROIs for Synaptophysin 1 and pTau (i.e., Tau-positive and DAPI-negative). Next, the average pixel intensity of every ROI was calculated and imported into Excel spreadsheet, in which it was subtracted by the average pixel intensity of the background ROIs from the same image. All resulting intensity values were randomized and 111 of them from every brain slice were imported into Prism (10.2.3, GraphPad) for statistical analyses.

***Statistics***

In every electrophysiological test, we used data from all 9 neurons in nine brain slices (one per slice) prepared from three mice (three slice per mouse) for every treatment (i.e., DMSO and three OGAi). We first used Welch’s ANOVA test to determine if there were any significance differences across all four treatments. Then, we used Kolmogorov-Smimov test to compare every OGAi treatment with DMSO control. The inset violin plot in Figure 1C represented the average value for every neuron for 60-90 seconds whereas the main plot represented the averages and the standard deviations of data points from all 9 neurons at every time point. For immunofluorescence comparison, we used all 9 slices and performed Welch’s ANOVA test followed by Kolmogorov-Smimov test to compare individual OGAi with DMSO control. In both figures, *, 0.05 > *p* ≥ 0.01; **, 0.01 > *p* ≥ 0.005; ***, 0.005 > *p* ≥ 0.001, ****, *p* < 0.001.
